# Continuous single cell transcriptome dynamics reveal a default vascular smooth muscle fate of FLK1 mesoderm

**DOI:** 10.1101/374629

**Authors:** Haiyong Zhao, Kyunghee Choi

## Abstract

Blood and endothelial cells arise from hemangiogenic progenitors that are specified from FLK1-expressing mesoderm by the transcription factor ETV2. FLK1 mesoderm also contributes to other tissues, including vascular smooth muscle (VSM) and cardiomyocytes. However, the developmental process of FLK1 mesoderm generation and its allocation to various cell fates remain obscure. Recent single cell RNA-sequencing (scRNA-seq) studies of early stages of embryos, or in vitro differentiated human embryonic stem (ES) cells have provided unprecedented information on the spatiotemporal resolution of cells in embryogenesis. These snapshots nonetheless offer insufficient information on dynamic developmental processes due to inadvertently missing intermediate states and unavoidable batch effects. Here we performed scRNA-seq of in vitro differentiated ES cells as well as extraembryonic yolk sac cells, which contain the very first arising hemangiogenic and VSM lineages, to capture the continuous developmental process leading to hemangiogenesis. We found that hemangiogenic progenitors from ES cells develop through intermediate gastrulation stages, which were gradually specified by ‘relay’-like highly overlapping transcription factor modules. Unexpectedly, VSM and hemangiogenic lineages share the closest transcriptional program. Moreover, transcriptional program of the *Flk1* mesoderm was maintained in the VSM lineage, suggesting the VSM lineage may be the default pathway of FLK1 mesoderm. We also identified cell adhesion signals possibly contributing to ETV2-mediated activation of the hemangiogenic program. This continuous transcriptome map will facilitate both basic and applied studies of mesoderm and its derivatives.

## Introduction

Previous studies have established that blood and endothelial cells differentiate from FLK1-expressing mesoderm^1^. FLK1 mesoderm can also contribute to vascular smooth muscle (VSM), cardiac lineages, and paraxial mesoderm tissues^2, 3^. However, high plasticity, heterogeneity, and rapid differentiation of early embryo cells have hampered investigations on the lineage relationships among these cell types, the mechanisms underlying the lineage restriction steps from pluripotent cells to FLK1 mesoderm, and the following commitment to different downstream lineages. Recent progress in scRNA-seq technology has unprecedentedly advanced our understanding of the complex cell constitution at discrete time points^4–7^ or in sorted cell populations^8, 9^ in early embryo development or in vitro differentiated ES cells^10, 11^. However, none of these studies have provided a complete process from pluripotent cells to FLK1 mesoderm due to missing intermediate states or batch effects. Therefore, there is still a gap in our current knowledge on the continuous sequence of FLK1 mesoderm development and its segregation into the hemangiogenic lineage.

Mouse ES cells in culture differentiate in an asynchronous manner giving rise to embryoid bodies (EB). Both undifferentiated pluripotent cells and differentiated progenies^12, 13^ coexist in mouse EBs of early stages of differentiation. We reasoned that cells in the intermediate states also exist in the same culture, representing the full spectrum of differentiation, which would allow a unique opportunity to delineate a continuous differentiation process without batch effects. By taking advantage of the asynchronous nature of the differentiation of mouse ES cells and the culture conditions that favor hemangiogenic lineage differentiation, we performed scRNA-seq of EB cells to elucidate a continuous developmental path from the pluripotent state to FLK1 mesoderm, and to the hemangiogenic lineage. We found that the cardiac lineage branched out immediately after nascent mesoderm specification.

Intermediate states positioned in-between ES and FLK1 mesoderm were distinguished by a few temporally overlapping transcription factor modules, suggesting a plastic and combinational regulatory mechanism in *Flk1* mesoderm development. Notably, expression of most of the characteristic genes of *Flk1* mesoderm continuously increased in the VSM branch, implying a ‘default’ route. Our data suggested that ETV2 turns on the hemangiogenic program from FLK1 mesoderm by restraining the VSM fate. We further identified cell adhesion-SRC kinase signaling as a potential regulator of ETV2-mediated hemangiogenic lineage specification. These studies provide a comprehensive description of mesoderm and hematopoietic/endothelial cell lineage development at the single cell level.

## Results

### EBs contain undifferentiated pluripotent cells and FLK1 mesoderm tissues

We performed scRNA-seq using day 4 EB cells in hemangiogenic differentiation. After filtering out low-quality cells, 1848 cells were kept for further analysis, which had 4373 genes/27272 unique mRNAs detected per cell in average (Figure 1A). Day 4 EB cells were grouped into 18 populations (Figure 1B, supplemental Figure 1A). 936 genes were differentially upregulated across the populations (supplemental Table 1). t-Stochastic Neighbor Embedding (t-SNE) indicated that most clusters did not form a distinct, well-segregated population (Figure 1B). Instead, distinct populations branched out gradually from the trunk, underpinning continuous developmental processes. We assigned different fates to the branches according to the expression of related marker genes (supplemental Table 1). SPRING^14^, a different tool for high dimension scRNA-seq data visualization, produced a similar cell clustering pattern (supplemental Figure 1B). High expression of *Sox2*, *Pou5f1/Oct4*, *Nanog* and *Zfp42/Rex1* highlights the maintenance of undifferentiated naïve pluripotent cells in the EBs (Figure 1C, 1D). FLK1 mesoderm and the hemangiogenic lineage, as marked by high expression of *Flk1*/*Kdr* and *Etv2*, and upregulation of the ETV2 target genes *Tal1* and *Gata2*, clustered at the other end of the ES cell population (Figure 1C, 1D). Expression of related genes showed a gradual shift in intermediate clusters from the pluripotent ES population to the *Flk1* mesoderm branch, revealing continuous intermediate states from ES to FLK1 mesoderm that are accessible in EBs.

**Figure 1.**
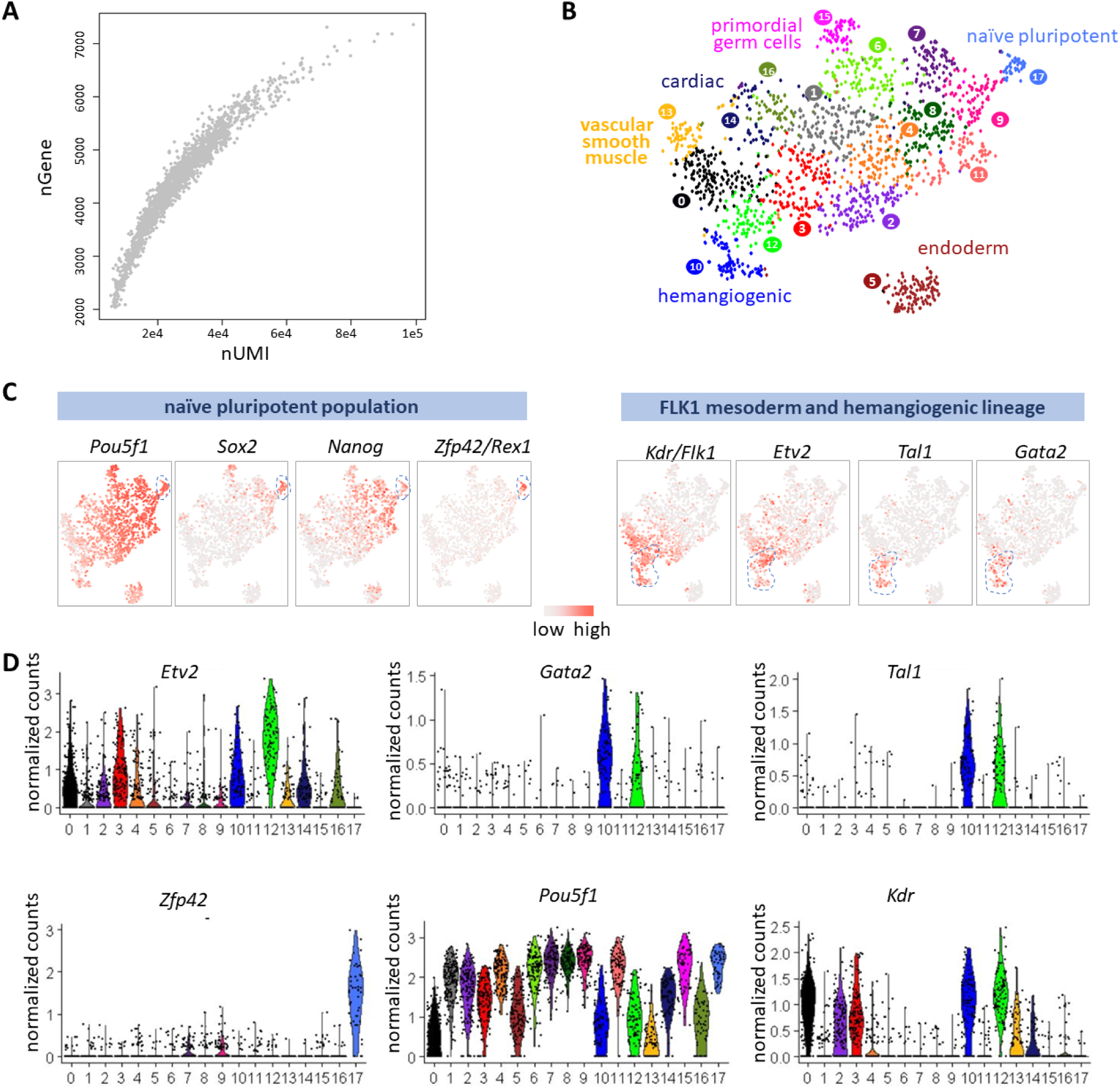
scRNA-seq of EBs containing undifferentiated pluripotent cells and differentiated cells. (A) Distribution of genes and transcripts detected in individual cells. (B) t-SNE projection of all cells. Cluster numbers is corresponding to those in supplemental Figure 1A. Potential lineage branches were annotated according to specific marker gene expression patterns. (C-D) Expression of the indicated marker genes for specific cell states/lineage fates in the t-SNE.

### scRNA-seq captured multiple intermediate stages from naïve pluripotency to FLK1 derivatives in EBs

To further determine developmental relationships among the clusters in-between the ES and *Flk1* hemangiogenic populations, we first examined the dynamics of the loss of pluripotency and the bifurcation of mesoderm and endoderm fates. Pluripotency is maintained in two states: the ‘naïve’ and the ‘primed’^15^. The former represents the primitive phase and highly expresses *Zfp42/Rex1*; while the latter loses *Zfp42* expression and harbors an unstable pluripotency gene network, which marks a nascent stage of gastrulation/mesendoderm ^15^. We picked out clusters which are located in the endoderm path (4, 5, 8, 9, 11 and 17 in Figure 1B), and re-clustered the cells based on 163 most differentially expressed genes among these cells (Figure 2A). A narrow “path” from Utf1-expressing pluripotent population leads to the *Sox17*-expressing endoderm (red dash line-surrounded region), next to the sprouting *Mesp1-*expressing nascent mesoderm. We next ordered cells in the endoderm path into a pseudo-time developmental line and examined the kinetics of 163 most differentially expressed genes (Figure 2B-2D, supplemental Figure 2C). At the beginning of the path, we observed an exit from the naïve pluripotency (Figure 2C, flanking the blue dash line).

**Figure 2.**
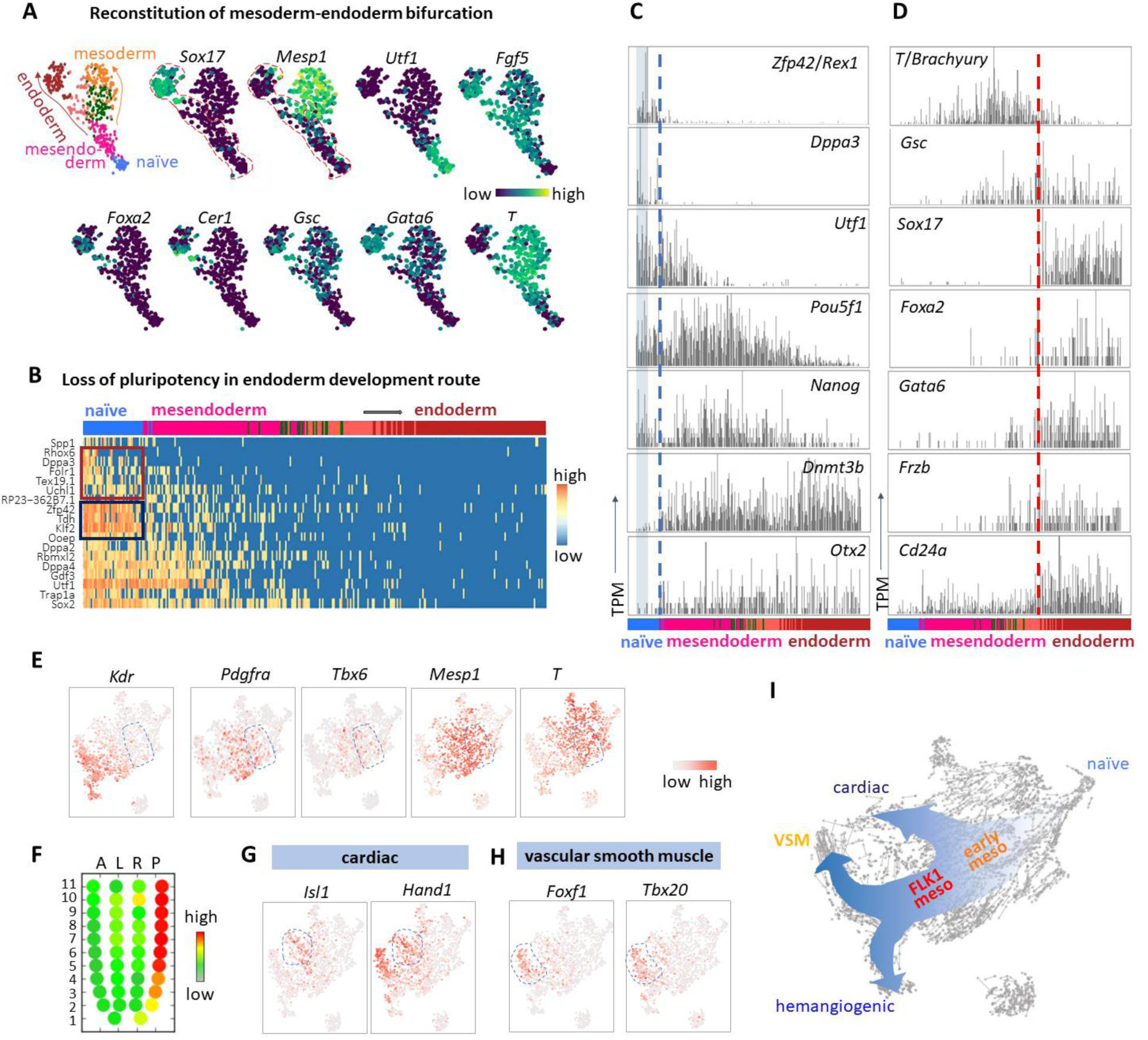
scRNA-seq captured continuous intermediates from naïve pluripotency to FLK1 derivatives. (A) Re-clustering of populations in exit from naïve pluripotency and bifurcation of endoderm and mesoderm. Clusters 4, 5, 8, 9, 11 and 17 in Figure 1B were picked to re-cluster based on Principle Component Analysis. The first two components are shown. The clusters are in the same colors as they are in Figure 1B. “naïve”, naïve pluripotency. (B) Cells in the endoderm differentiation route (red dash line-surrounded region) were ordered into pseudo-time line. Arrows indicate the differentiation direction. Heatmap of naive pluripotent cell-enriched genes are shown. Brown box encircles genes heterogeneously expressed in the naïve stage; blue box marks the constantly expressed naïve stage-specific genes. (C) Barplot of expression of selected marker genes along the endoderm pseudotime line. Shaded region covers cells maintaining high *Dppa3* expression in the naïve population. Blue dash line marks the exit point from naïve pluripotency. (D) Barplot of expression of marker genes along the pseudotime line of endoderm differentiation. Red dash line marks the point, where endoderm fate is committed. (E) Expression of indicated genes in early mesoderm cells. (F) Zipcode mapping of early mesoderm cells (cluster 4 in Figure 1B. Mean values of each genes in the cluster were used for the mapping) to E7.0 mouse embryo. (G-H) Expression of indicated genes. (I) scheme of development routes of indicated mesoderm lineages.

Unexpectedly, expression of a small group of genes (represented by *Dppa3*, brown box in Figure 2B, and in Figure 2C) already displayed strong heterogeneity in the *Zfp42*-expressing population. Gradual loss of the expression of these genes preceded the exit from naïve pluripotency. This may suggest an extra layer of guarding mechanisms of pluripotency beyond the naïve circuit (as represented by the genes in the blue box in Figure 2B, which have relatively constant expression in the naïve pluripotency stage). Primitive streak markers such as *T* and *Fgf5* were up regulated in cells exiting from the naïve pluripotent state and entering the mesendoderm stage (Figure 2A). Shortly after, the endoderm and mesoderm lineages bifurcated. *Fgf5* expression became more enriched in endoderm, while *T* expression was exclusively enriched in mesoderm (Figure 2A). *Gata6* and *Gsc* were expressed in both mesoderm and endoderm, although they were expressed more extensively in the latter (Figure 2A). The majority of cells in the endoderm population were separated from the trunk, with very few *Foxa2*-high expressing cells in the intermediate state. When we lowered down the stringency and included more variable genes to cluster the cells, the connection between endoderm and the mesendoderm cluster becomes more obvious (supplemental Figure 2A-B). These results might reflect that the core endoderm genes form a strong self-enforcing circuit, rendering the intermediate specification state relatively unstable.

The immediate upstream population of *Flk1*^+^ mesoderm expressed *Pdgfra* and *Mesp1* (Figure 2E, blue dash line-marked region). These cells also expressed low levels of *Tbx6*, implying that they are to some extent ‘primed’ for paraxial mesoderm, which generates tissues like skeletal muscle and cartilage. As such, they might represent an early ancestral state for both FLK1 lateral plate mesoderm and paraxial mesoderm. Consistently, cells in this region have an expression pattern similar to the posterior primitive streak of mouse embryo at E7.0, where nascent mesoderm emerges (Figure 2F). Furthermore, these cells were transcriptionally similar to *Mesp1*-labeled early mesoderm cell population in mouse E6.75 embryos in Lescroart et al’s recent scRNA-seq data^8^. This embryonic population of cells still expressed high level of the pluripotency marker *Pou5f1*, already expressed high levels of *Mesp1* and *Pdgfra*, while expressed low levels of *Tbx6 and* has not upregulated *Kdr* yet (cluster 1 in supplemental Figure 2D). The cardiac lineage branched out mainly from the *Pdgfrα*^+^*Flk1*^-^ early mesoderm population, while a small group of *Flk1* mesoderm also expressed the cardiac transcription factor *Isl1* (Figure 2G). Notably, VSM and hemangiogenic lineages directly bifurcated from a *Flk1*-expressing population, suggesting a common origination (Figure 2H). RNA velocity estimation of the variable genes, based on the abundance of unspliced and spliced mRNAs^16^, revealed persistent overall cell state dynamics from naive pluripotency to differentiation branches (supplemental Figure 2E). These results suggested a continuous process from pluripotency to *Flk1* mesoderm and its derivate tissues (Figure 2I).

### Cells corresponding to different development stages proliferate equally

It was possible that cells in alternative differentiation paths might have different proliferation rate. The expression pattern of typical S-phase marker genes showed no obvious bias in different cell populations (Figure 3A, supplemental Figure 3A). Using the cell cycle analysis tool in the Seurat package^17^, we assigned each cell to a specific phase in the cell cycle and found that the majority of cells in the mesoderm developmental route proliferated robustly (Figure 3B, supplemental Figure 3B). Analysis of cell proliferation using the violet proliferation dye 450 (VPD450) also confirmed that EB cells with different levels of differentiation (indicated by reporter of the hemangiogenic marker gene *Etv2*) proliferated similarly in our system (Figure 3C, top). As a control, ERK inhibitor PD0325901-mediated differentiation block and cell cycle delay was observed in the VPD450 dilution assay (Figure 3C, bottom).

**Figure 3.**
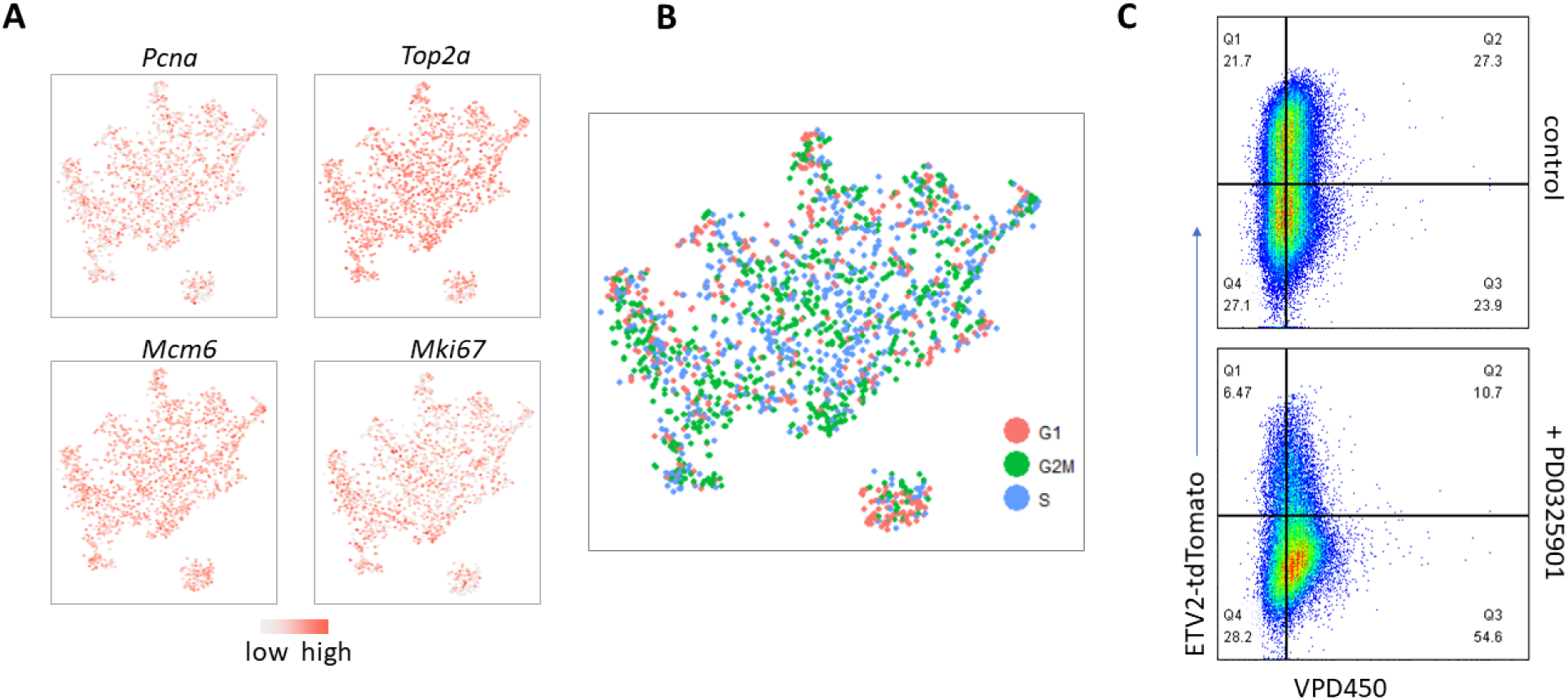
Cells corresponding to different development stages proliferate equally. (A) Expression of indicated cell cycle genes. (B) Assigned cell cycle phases of each cells. (C) Proliferation of cells in EBs treated with or without 2μM of ERK inhibitor PD0325901 from day 3 was compared based on dilution of the Dye VPD450. Dye dilution and the *Etv2*-tdTomato reporter gene expression were examined 24 hours later using flow cytometry.

### Transcriptome dynamics in hemangiogenic lineage development

Next, we retrieved a continuous process of hemangiogenic lineage development by manually choosing the populations corresponding to the shortest path from the naïve pluripotent state to the hemangiogenic branch and reconstructed a fine developmental route (Figure 4A). A group of *Flk1*-expressing cells achieved the highest level of *Etv2* expression, followed by dramatic up regulation of the ETV2 target genes, *Tal1*(*Scl*) and *Gata2*, confirming that the threshold-level of *Etv2* expression is necessary for hemangiogenic lineage specification^18^ (Figure 4A). We ordered the cells into a 1-D pseudo-time line and assigned the populations to specific stages according to the dynamics of marker gene expression (Figure 4B). 68 most variable transcription regulation-related factors were clustered based on their expression along the pseudo-time line (Figure 4C). They aggregated into a few obvious modules covering different developmental stages (blue dash line boxed regions in the heatmap).

**Figure 4.**
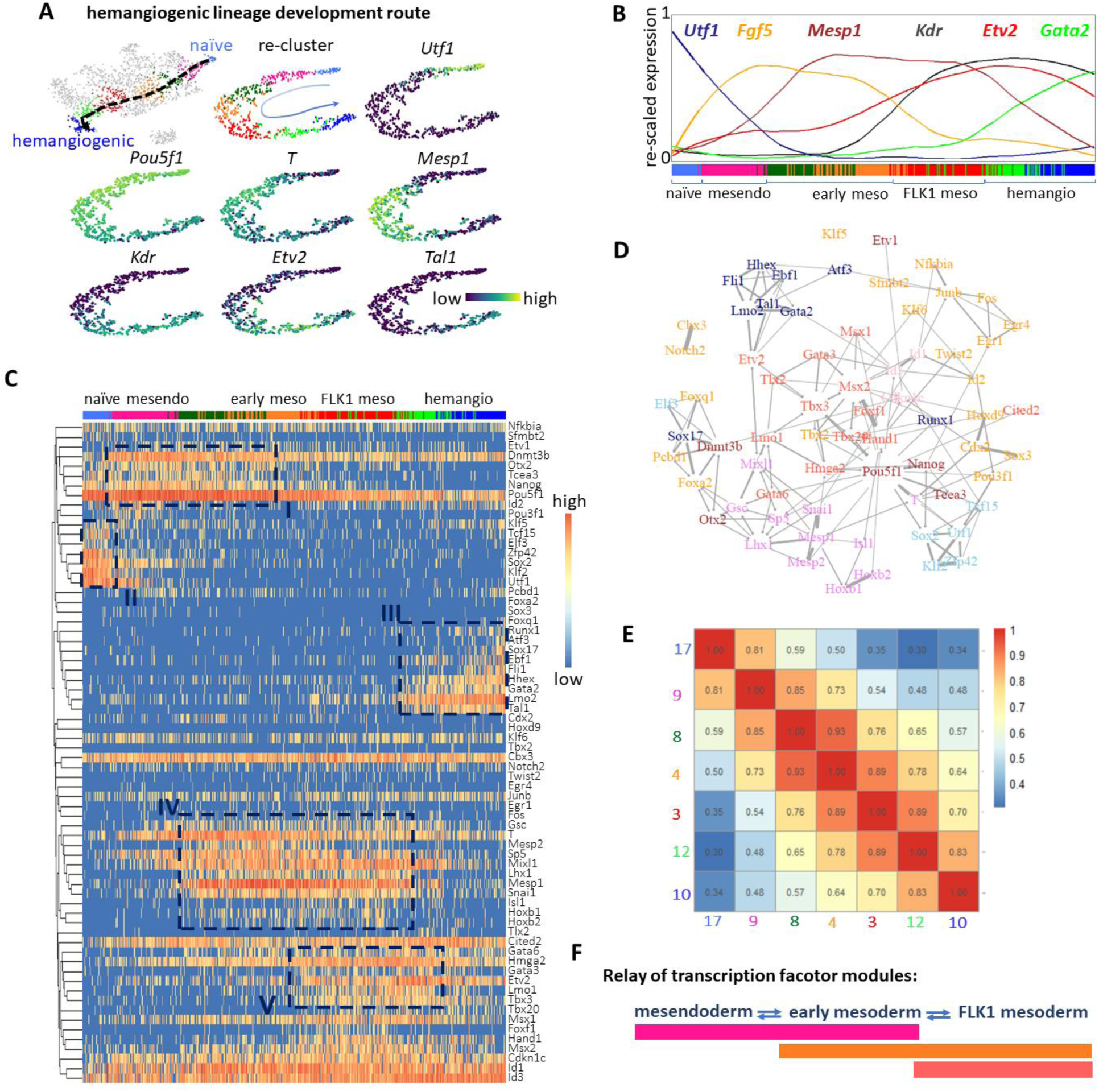
Underlying transcription factor modules in hemangiogenic lineage development. (A) Re-clustering of populations in the route of hemangiogenic lineage development. Clusters 3, 4, 8, 9, 10, 12 and 17 in Figure 1B were picked to re-cluster. The first two components of PCA are shown for re-clustered cells. The clusters are in the same colors as they are in Figure 1B. Arrows indicate the differentiation direction. Expression of the representative marker genes in the re-clustered populations is shown. (B) The re-clustered cells are ordered into a pseudo-time line of differentiation. Expression dynamics of representative marker genes along the pseudo-time line is shown. Gene expression data was smoothened. The expression curves and the gene names are in same colors. At the bottom, color bars indicate the cells’ original identities as in Figure 1B. Cell states were annotated: “naïve, naïve pluripotent state; “mesendo”, mesendoderm; “early meso”, early mesoderm; “FLK1 meso”, FLK1 mesoderm; “hemangio”, hemangiogenic lineage. (C) Dynamics of transcription-related factors from the 427 most variable genes is shown along the pseudo-time line of hemangiogenic lineage development. The genes are hierarchically clustered based on Pearson’s correlation. Boxes in a yellow dash-line indicate the obvious stage-specific gene modules. (D) Gene regulatory network of the trancription-related factors shown in Figure 4C. Only the top250 gene-gene links were shown. The width of the edges indicated the weight of the links. (E) Gene expression similarities among individual clusters along the hemangiogenic development route. The mean values of gene expression of cells in individual clusters were used to calculate the Pearson’s correlation scores. (F) Scheme of the ‘relay’-like transcription factor modules underlying FLK1 development.

Notably, the naïve pluripotency module (box II, Figure 4C) and the hemangiogenic module (box III) were relatively independent, while each of the intermediate gastrulation modules were shared by adjacent stages (box I, IV, and V, Figure 4C). The gene network constructed based on the gene co-expression also highlighted the collaboration among transcription factors of individual modules (Figure 4D). This stepwise combinatorial usage of different regulatory circuits fits the diverse lineage specifications in gastrulation. Consistently, the intermediate populations were less committed, largely interchangeable^19^, and more similar to each other in transcriptome (Figure 4E, supplemental Figure 4A, 4B). These results suggested that the intermediate states constitute a ‘relay’-like multistep specification process (Figure 4F). Exit from the naive pluripotency and activation of the hemangiogenic program might be the two critical transitions in hemangiogenic lineage development.

### Cell adhesion signaling potentiates ETV2-mediated hemangiogenic fate specification

Due to intracellular noises and microenvironmental influences, gene expression within individual cells keeps fluctuating. scRNA-seq analysis offers an opportunity to utilize this luxuriant gene expression heterogeneity to assess a regulatory network. We previously reported that *Etv2* expression above a threshold-level is essential to activate hemangiogenic genes^18^. We confirmed this at a single cell endogenous mRNA level (Figure 5A). We noticed that the kinetics of upregulation of ETV2 target genes soon became extremely steep after *Etv2* expression reached to a specific level, suggesting an *Etv2* mediated sensitive “switch” mechanism in activating its downstream target genes. We explored potential ETV2 cooperating genes for triggering this sensitive switch by examining potential upstream regulators of *Tal1* and *Lmo2*, two ETV2 direct target genes that are critical for establishing the hemangiogenic lineage^20, 21^. First, we assumed that if more gene A-expressing cells simultaneously expressed gene B, it is likely that gene A expression depends on gene B (Figure 5B, see methods for more details). From the population where ETV2 starts to activate hemangiogenic genes (population 12 in Figure 1B), we assessed the possibility of a given gene being required for *Tal1* or *Lmo2*’s expression (Figure 5C, 5D). *Etv2* and *Kdr/Flk1* were at the top of the list of genes required for *Tal1* or *Lmo2* expression (Figure 5C, 5D, supplemental Table 2), consistent with the finding that VEGF-FLK1 signaling ensures *Etv2* threshold expression to activate hemangiogenic genes^18^. The comparison of the top 100 genes required for *Tal1* expression and the top 100 required for *Lmo2* expression generated 66 in common, of which 22 were annotated to cell adhesion/MAPK-related signaling (supplemental Figure 5A, highlighted in supplemental Table 2). The majority of these 22 genes were already highly expressed in populations preceding the hemangiogenic stage (supplemental Figure 5B). To test if cell adhesion signaling plays a role in ETV2-mediated hemangiogenic gene activation, we utilized differentiation of doxycycline (DOX)-inducible *Etv2*-expressing ES cell line^22^. Since extracellular matrix/cell adhesion signaling converges on the SRC kinase^22^, we added SRC inhibitor PP2 to day 3 EBs, while simultaneously inducing exogenous *Etv2* expression. After 24 hours, as expected, DOX-induced *Etv2* expression greatly enhanced the FLK1^+^PDGFRα^-^ population that enrich hemangiogenic lineage cells ^22^, while inhibition of SRC using PP2 was sufficient to reduce *Etv2* overexpression-induced hemangiogenic lineage skewing (supplemental Figure 5C). Given that cell adhesion is necessary for gastrulation as well^23^, it was possible that PP2 affected earlier stages rather than directly affecting ETV2’s function. To minimize this possibility, we induced ETV2-mCherry fusion protein expression ectopically in undifferentiated ES cells, where endogenous *Etv2* level is ignorable^18^ and exogenous *Etv2* can induce *Flk1* expression.

*Flk1* is also a target of ETV2 and reciprocal activation between ETV2 and FLK1 signaling is an important mechanism for hemangiogenic fate determination^18^. Exogenous ETV2 induced more than 8% of FLK1-positive cells in undifferentiated ES cells, while this ratio was significantly reduced by PP2 treatment (Figure 5E). Unexpectedly, PP2 treatment also dramatically reduced ETV2 protein level, with only a limited number of cells achieving threshold expression of *Etv2* (Figure 5E). To exclude the possibilities that PP2-mediated SRC inhibition affected DOX-induction system or general gene expression processes, we tested DOX-induced expression of another transcription factor, EOMES, and the constitutively active promoter *EF1α*-driven expression of the red fluorescent protein mCherry.

**Figure 5.**
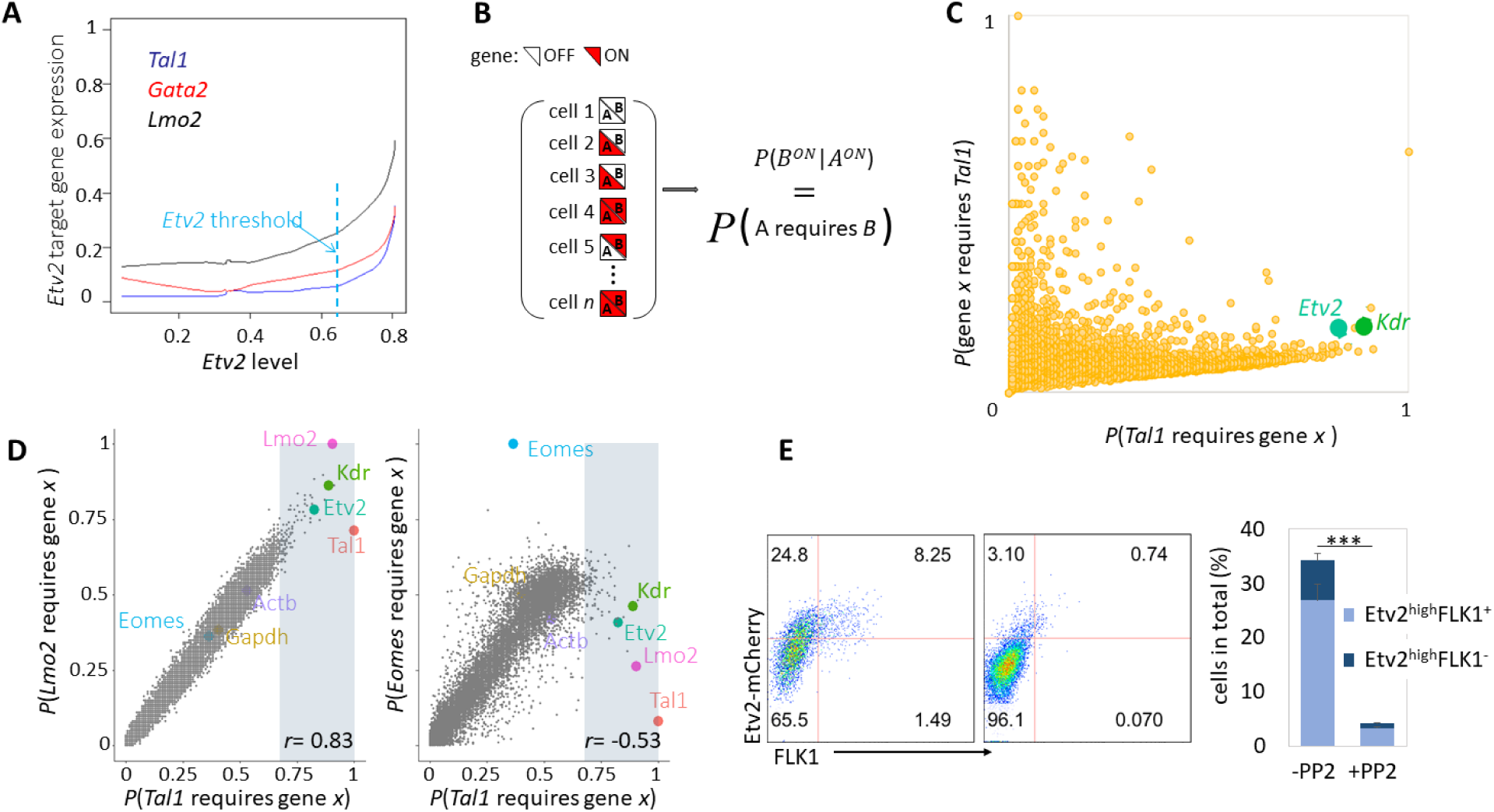
ETV2 triggers a sensitive switch initiating hemangiogenic program, depending on cell adhesion signaling. (A) ETV2 target genes sensitively respond to *Etv2* dosage above the threshold. The input data was from smoothed and re-scaled expression of related genes along the pseudo-time line of hemangiogenic lineage development. Datasets only before *Etv2* achieves its highest level were displayed, because after that *Etv2* starts to be down regulated. (B) Scheme for identification of potential regulators of ETV2-mediated hemangiogenic fate specification. If gene B is required for gene A expression, then in a cell where gene A is expressed gene B should also be expressed. In a population, more gene A expressing-cells have gene B expression, it is more likely that gene B is required for gene A expression. (C) All detected genes’ relationship with *Tal1*. x-axis is the possibility of *Tal1* expression requires a given gene *x* in group 12 in Figure 1C, where *Etv2* achieves threshold expression and starts to activate hemangiongenic genes. y-axis is the possibility of *Tal1* in all cells in return is required for a specific gene’s expression. *Etv2* and *Kdr/Flk1*, two known upstream regulators of *Tal1*, are labeled in the plot. (D) Comparison of genes required for *Tal1* expression to that required for *Lmo2* or *Eomes*. *Eomes* is a FLK1 mesoderm-expressing transcription factor but not reported regulating hemangiogenic genes. For the top 100 genes that most tightly required for *Tal1* expression in the cluster 12 (dots in shadow regions), the Pearson’s correlation coefficient (*r*) between x-axis and y-axis are shown respectively. Genes of interest are shown in color. House keeping genes *Actb* (beta-actin) and *Gapdh* are shown as control. (E) A2lox ES cells with DOX-inducible ETV2-mCherry were transferred to feeder-free condition and cultured in the presence of 0.5μg/mL of DOX, simultaneously treated with or without PP2 (5μM) for 24 hours. Cells were then analyzed for FLK1 and mCherry levels using flow cytometry. The horizontal red lines mark ETV2 threshold. The statistics summary is shown on the right. ***, P<0.001.

We found that DOX-induced EOMES protein level only mildly decreased in the presence of PP2, while *EF1α*-driven mCherry expression was insensitive to this treatment (supplemental Figure 5D). These results suggest that cell adhesion signaling contributes to ETV2-mediated hemangiogenic fate specification via regulation of ETV2 protein production.

### FLK1 mesoderm is fated to the vascular smooth muscle (VSM) lineage, while ETV2 skews it into the hemangiogenic fate

Based on the observation of direct bifurcations of VSM and hemangiogenic lineages from FLK1 mesoderm (Figure 2I), we analyzed the cell populations corresponding to the developmental route of the VSM lineage (Figure 6A, supplemental Figure 6A), and examined expression of the 427 most varied genes along the pseudo-time of differentiation (Figure 6B). Many of the variable genes aggregated as modules. The modules corresponding to mesendoderm, nascent mesoderm, and *Flk1-*expressing mesoderm stages displayed largely overlapping ‘relay’-like patterns. Three groups of genes were most extensively expressed in the VSM branch (Figure 6B, gene clusters a, b and c in the black dash line boxed region). Of the three groups of genes, group b and c were already up regulated prominently in FLK1 mesoderm, with the cluster c being up regulated even earlier from the mesendoderm stage. The transcription factors *Hand1*, *Tbx20*, and *Foxf1* were among the gene cluster b (Figure 6B, 6C). *Hand1* and *Foxf1* have been reported to be important for VSM development^24, 25^.

**Figure 6.**
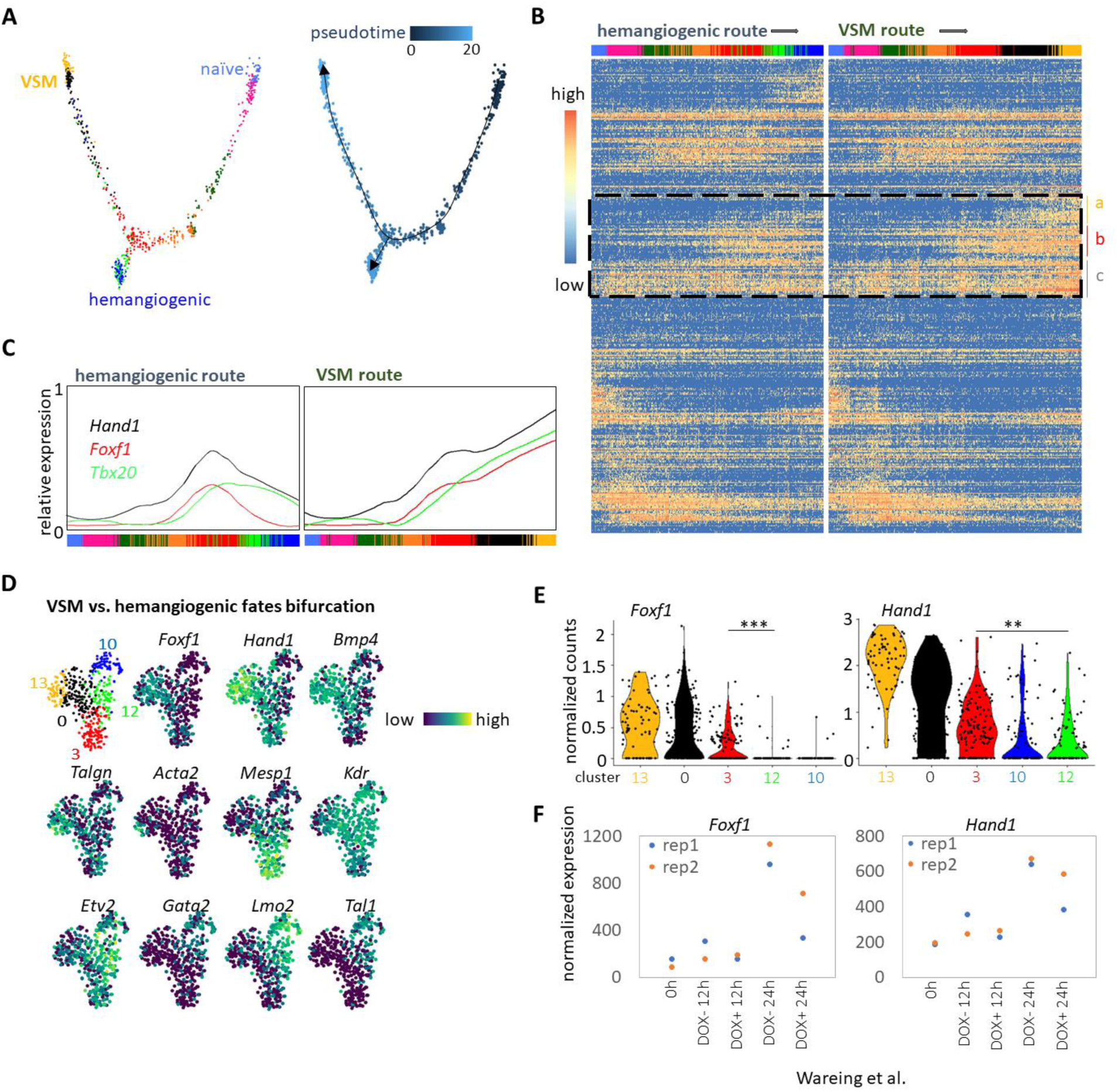
‘Default’ vascular smooth muscle fate of FLK1 mesoderm. (A) Left, reclustering of populations in the routes of VSM and hemangiogenic lineage development. Clusters 0, 3, 4, 8, 9, 10, 12, 13 and 17 in Figure 1B were picked, and the first two components of PCA are shown. Right, pseudotime analysis of the reclustered cells. (B) The comparison of the most variable 427 genes’ dynamics along the VSM or hemangiogenic routes is shown. The genes were clustered based on Pearson’s correlation. Color bars indicate cells’ original identities. “a”, “b”, and “c”, shown in a yellow dash-box, mark three groups of genes that are mostly enriched in the VSM branch. (C) Expression dynamics of indicated genes along the pseudo-time lines of hemangiogenic and VSM lineage development is shown. The expression values were smoothened. The colors of curves and gene names are consistent. (D) Clusters 0, 3, 10, 12 and 13, corresponding to the VSM/hemangiogenic lineage bifurcation fork in Figure 1B were picked and reclustered. Cells are colored according to the expression levels of indicated genes. (E) Violin plots comparing indicated genes’ expression in different populations. **, P<0.01; ***, P<0.001. (F) Important VSM transcription factors respond to Etv2 overexpression. Day 2.5 *Etv2*-knockout EBs were induced for exogenous *Etv2* expression using DOX. Two replicates of samples corresponding to DOX induction for 0 hour, 12 hours or 24 hours were collected for microarray analysis.

Expression of the genes in clusters b and c remained similar in the hemangiogenic branch. Only a small cluster of genes was exclusively up regulated after cells entered the VSM branch (gene cluster a). Importantly, all the VSM transcription factor genes, except *Tbx2*, were already extensively expressed in the FLK1 mesoderm and constituted a major characteristic of the FLK1 mesoderm (supplemental Figure 6B, in the black dashed box). These results suggest that VSM lineage might be the default fate of the FLK1 mesoderm.

We zoomed into the populations 0, 3, 10, 12 and 13 (as labeled in Figure 1B), corresponding to the bifurcation of hemangiogenic and VSM fates (Figure 6D). Expression of *Bmp4*, which can enhance VSM development^26^, was dramatically up regulated in the VSM branch (from population 0 (black) to population 13 (orange)), but not in population 12 (green), where *Etv2* initiates specification of the hemangiogenic fate (Figure 6D). *Hand1* and *Foxf1* were dramatically down regulated in population 12 compared to the upstream population 3, implying an active repression of them by *Etv2* (Figure 6E). Consistently, by analyzing Wareing and colleagues’ work^27^ we found that re-expression of *Etv2* in *Etv2*-knockout EBs inhibited expression of *Hand1* and *Foxf1* (Figure 6F). These results suggest that *Etv2* represses the default VSM program when specifying the hemangiogenic lineage from FLK1 mesoderm.

### FLK1 mesoderm characteristics are enriched in vascular smooth muscle tissue in vivo

To better understand the VSM program, we further analyzed the overall expression score of the group b genes in Figure 6B (listed in supplemental Table 3). The group b gene score reflected the dynamics of FLK1 mesoderm identity in day 4 EB data (supplemental Figure 7A). The cardiac lineage progenitors also had a high group b gene score; as such we termed the group b gene score as the lateral plate mesoderm score, ‘LPM-score’ (both FLK1 mesoderm and cardiac tissues arise from lateral plate mesoderm). If indeed *Flk1* mesoderm default fate is VSM, it was expected that *Flk1* mesoderm identities would persist in the VSM lineage. To determine if this is the case in vivo, we performed scRNA-seq of mouse extraembryonic yolk sac, a tissue that generates the first arising hemangiogenic and VSM lineages (Figure 7A). We found that the LPM-score was particularly high in the VSM tissue in mouse yolk sac (Figure 7B), exemplified by the coexpression of the smooth muscle actin gene *Acta2* and the transcription factor *Foxf1* and *Hand1*, and the absence of the expression of hematopoietic and endothelial transcription fator *Tal1/Scl* (Figure 7C, supplemental Figure 7B). Ibarra-Soria et al. recently reported scRNA-seq of E8.25 mouse embyros^5^. A group of cells in their data, which they assigned as ‘amnion’, had high LPM-score and expressed VSM markers (Figure 7D-F, supplemental Figure 7C-D). Presumably, this ‘amnion’ population represents embryonic VSM. Consistently, the 72 marker genes shared by day 4 EB VSM, yolk sac VSM, and ‘amnion’ were enriched with smooth muscle function annotation (Figure 7G-H). Collectively, the *Flk1* mesoderm characteristics persist in embryonic and extraembryonic VSM tissues.

**Figure 7.**
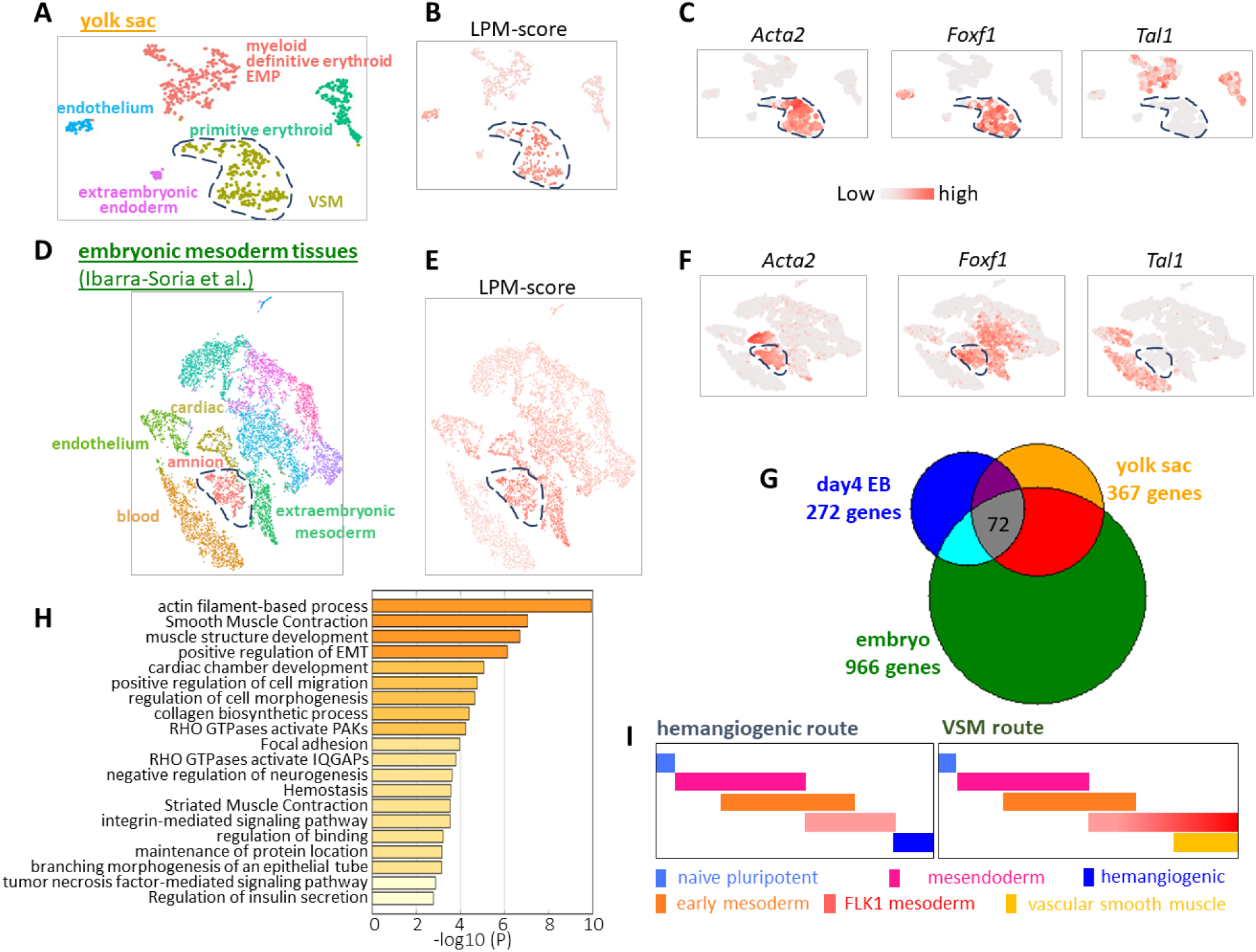
Characteristics of FLK1 mesoderm are enriched in vascular smooth muscle tissue in vivo. (A) t-SNE plot of scRNA-seq analysis of yolk sac. (B) LPM-score gene expression in E8.5 yolk sac cells. (C) Expression of indicated genes in E8.5 yolk sac. (D) Re-analysis of Ibarra-Soria et al.’s^5^ scRNA-seq of E8.25 embryo. Only mesoderm tissues were shown. (E-F) Expression of indicated genes in Ibarra-Soria et al.’s mesoderm population. (G) Comparison of VSM marker genes from indicated datasets. (H) Annotation of the shared 72 VSM genes by the three different datasets. (I) Model of gene module dynamics along VSM and hemangiogenic lineage development.

## Discussion

It has been reported that single FLK1*^+^* mesoderm cells can form both VSM and endothelial cells in culture^2^. Moreover, sorted FLK1*^+^* cells are able to form cardiomyocytes and even skeletal muscles^19, 22^. However, the lineage relationships among cardiac, vascular smooth muscle, skeletal muscle, and hemangiogenic tissues concerning FLK1 mesoderm allocation remain elusive. Our single cell transcriptome profiling of EB cells revealed a continuous differentiation trajectory. We showed that the cardiac branch mainly segregates out from *Pdgfrα^+^Flk1^-^* nascent mesoderm. Unexpectedly, VSM and hemangiogenic lineages have the closest lineage relationship and directly bifurcate from a common group of *Flk1*-expressing cells. Notably, most FLK1 mesoderm stage markers were further upregulated in the VSM branch, suggesting that VSM might be a default fate of the *Flk1*-expressing mesoderm. We propose that *Etv2* drives the branching of the hemangiogenic lineage from the VSM fate.

Expression of cell adhesion signaling coincides with ETV2 target gene activation for hemangiogenic lineage specification. Consistently, inhibition of SRC, a kinase important for cell adhesion signaling, was sufficient to repress hemangiogenic lineage skewing induced by *Etv2* overexpression. Meanwhile, critical VSM transcription factors, *Hand1* and *Foxf1*, become actively down regulated. Notably, neither *Hand1* nor *Foxf1* are direct ETV2 target genes^21^. It will be critical in the future to explore how this repression is achieved.

Hemangiogenic lineage development can be clearly annotated by a series of transcription factor modules, therefore providing strong molecular support for current definition of early embryo developmental stages (Figure 4C). Cells in the gastrulation stages, after exiting pluripotency and before entering the hemangiogenic state, are overall similar to each other in transcriptome, and the underlying transcription factor modules show a ‘relay’-like highly overlapping pattern. Consistently, FLK1^-^PDGFRα^-^, FLK1^-^PDGFRα^+^ and FLK1^+^PDGFRα^+^ populations in early differentiation of ES cells are reversible/interchangeable ^19^. These results suggest that exit from the naïve pluripotency state and activation of the hemangiogenic program might be the two rate-limiting steps in hemangiogenic lineage development, and that the intermediate gastrulating stages are plastic and adapted for stepwise specification to multiple lineage fates through combinational usage of limited regulatory circuits (Figure 7I).

In summary, our work described a continuous process of a mesoderm lineage tree leading to hemangiogenic and VSM tissue emergence and underlying molecular networks, thereby providing comprehensive information on early embryo and blood/blood vessel development. These findings are fundamental for the study of basic cell fate determination and gene network structures, and for designing more effective strategies to generate hematopoietic and endothelial cells for regenerative medicine.

## Methods

### Mouse ES cell culture and differentiation

Mouse ES cells were maintained and for EB differentiation in serum as previously reported ^18^. Briefly, ES cells were maintained on mouse embryo fibroblast (MEF) feeder cell layers in Dulbecco-modified Eagle medium containing 15% fetal bovine serum, 100 units/mL LIF, 1× MEM Non-Essential Amino Acids Solution (Gibco), 1× Glutamax^TM^ Supplement (Gibco), and 4.5 × 10^−4^ M 1-Thioglycerol (MTG, Sigma). For feeder-free culture, ES cells were maintained in gelatin-coated dish in Iscove’s modified Dulbecco medium (IMDM) with the same supplements as used for maintenance on feeder. For EB differentiation, ES cells were first transferred into feeder-free condition for 2 days, then single-cell suspensions were prepared, and 8,000 cells were added per mL to a differentiation medium of IMDM containing 15% differentiation-screened fetal calf serum, 1× Glutamax, 50 μg/mL ascorbic acid, and 4.5 × 10^−4^ M MTG on a bacteriological Petri dish. The SRC inhibitor PP2 (Sigma) and Doxycycline (Sigma) treatment were performed as indicated in the text.

### scRNA-seq

Yolk sacs were dissected from equal numbers of mouse E9.5 and E10.5 embryos, mixed together and dissociated with 0.25% collagenase at 37°C for 30 minutes.

Day 4 EBs were dissociated with Accutase solution (Sigma) for 5 minutes into single cells. The single cell suspension was washed with and re-suspended in PBS. Single cell suspension at 300 cells/μL in PBS were subjected to the Chromium 10X Genomics library construction and HiSeq2500 sequencing (The Genome Technology Access Center, Washington University in St. Louis). The sequenced reads were mapped to the GRCm38 assembly using Cell Range 2.0.1 (10x Genomics).

### scRNA-seq data QC

The scRNA-seq output was imported into Seurat 1.4 ^17^, and genes expressed in at least 3 cells were kept for analysis. Cells with more than 5% mitochondria reads or less than 2000 unique genes detected were filtered out. 1848 cells were kept for further analysis, which had 4373 genes/27272 unique mRNAs detected per cell in average. Gene expression counts were normalized and log transformed.

### scRNA-seq data visualization

According to the average expression and dispersion gene expression of each gene, we selected 427 most variable genes for further analysis of the remaining cells. nUMIs were re-scaled to regress the effects of detection depth of each cell and mitochondria reads out. The re-scaled data was run for principle components analysis to reduce dimensions. According to the PCA results the cells were clustered into 18 populations, and also used for t-SNE(t-distributed stochastic neighbor embedding) presentation. Cells in selected populations were imported into Monocle 2^28^ for re-clustering, PCA plotting and pseudo-time ordering.

For hierarchical clustering and heatmap plotting of selected genes or cells, the R package ‘pheatmap’ was used. All scRNA-seq data analyses were finished in R environment.

For cell cycle score and phase asignment with Seurat, the ‘CellCycleScoring’ function was invoked.

For LPM-score gene analysis, group b genes in Figure 6B were read and processed by invoking the ‘AddModuleScore’ function in Seurat.

For SPRING visualization, a matrix of expression counts of the 427 most variable genes vs. the filtered cells was uploaded (https://kleintools.hms.harvard.edu/tools/spring.html) per the guidence^14^. Cell cluster information from Seurat analysis was also loaded for viewing.

For RNA velocity estimation, the bam file in 10x genomics output data was first re-counted with the Velocyto counting pipeline in python according to the manual^16^. The generated loom file was loaded to velocyto.R. Only counts of the 427 most variable genes were used for RNA velocity estimation. Finally, coordinates from Seurat’s t-SNE analysis were used to embed the velocity results.

For gene regulatory network analysis, the R package GENIE3^29^ was used. To simplify the results, only transcription-related genes in the 427 most variable genes were used to calculate gene-gene link matrix, and only the top 250 weighted link were adopted for visulation. To view the network, the R package igraph was used.

For Zipcode mapping, mean expression of all the genes in chosen cells was uploaded to www.itranscriptome.org/ according to the developer’s guidence^30^.

### Identify genes required for ETV2-target gene activation

In cell cluster 12, the cells were subset into two groups, one with *Tal1* expression above the mean *Tal1* level across the whole population, and the other with *Tal1* below the mean level. In the first group, cells with gene *x* expression above its mean level, while in the second group, cells with gene *x* expression below its mean level, were counted seperately, and the sum of the two counts was divided by the total cell number of cluster 12. This ratio was considered as the possibility of gene *x* following *Tal1*. In fact, close gene-gene correlations can root from three possibilities, take *Tal1* as an example: 1) gene x lies upstream of *Tal1* and is required for its expression, 2) gene x is TAL1’s target, 3) both gene x and *Tal1* are ETV2’s target, thus respond to ETV2 in a similar way. However, in the population where expression of *Tal1* is just initiated, it is not likely that TAL1 activates its own target genes yet, therefore reduces possibility 2. We can refer to a gene’s expression pattern in the differentiation trajectory to test possibility 3. In fact, most of the genes tightly following *Tal1* and *Lmo2* were already extensively activated before ETV2 starts to function.

### Dye dilution assays

*Etv2*-tdTomato ES cells^18^ is dissociated into single cell suspension and incubated with 1μM of VPD450 (BD Horizon™) in PBS for 10 minutes at room temperature. Then the cells were aggregated for normal differentiation.

### Microarray data analysis

Wareing and colleagues’ microarray data^27^ was re-analyzed using the Expression Console software (Affymetrix).

### Gene functional annotation

Gene funcitonal annotation was performed using Metascape (http://metascape.org/gp/index.html)31.

### Flow cytometry

Single cells were incubated with α-mouse FLK1 (BioLegend) and α-mouse PDGFRα (BioLegend). Data were acquired on LSR-Fortessa flow cytometer (BDbiosciences) and analyzed using the FlowJo (Treestar) software.

### Statistical analysis

The results of flow cytometry were analyzed using Students’ t test. P < 0.05 was considered significant.

### Data availability

The data generated in this study can be downloaded from the NCBI Gene Expression Omnibus under accession number GSE130146.

## Acknowledgements

We thank members of the Choi lab for helpful discussion and support. The work was supported by NIH grant HL55337.

---

**Figure 1S.**
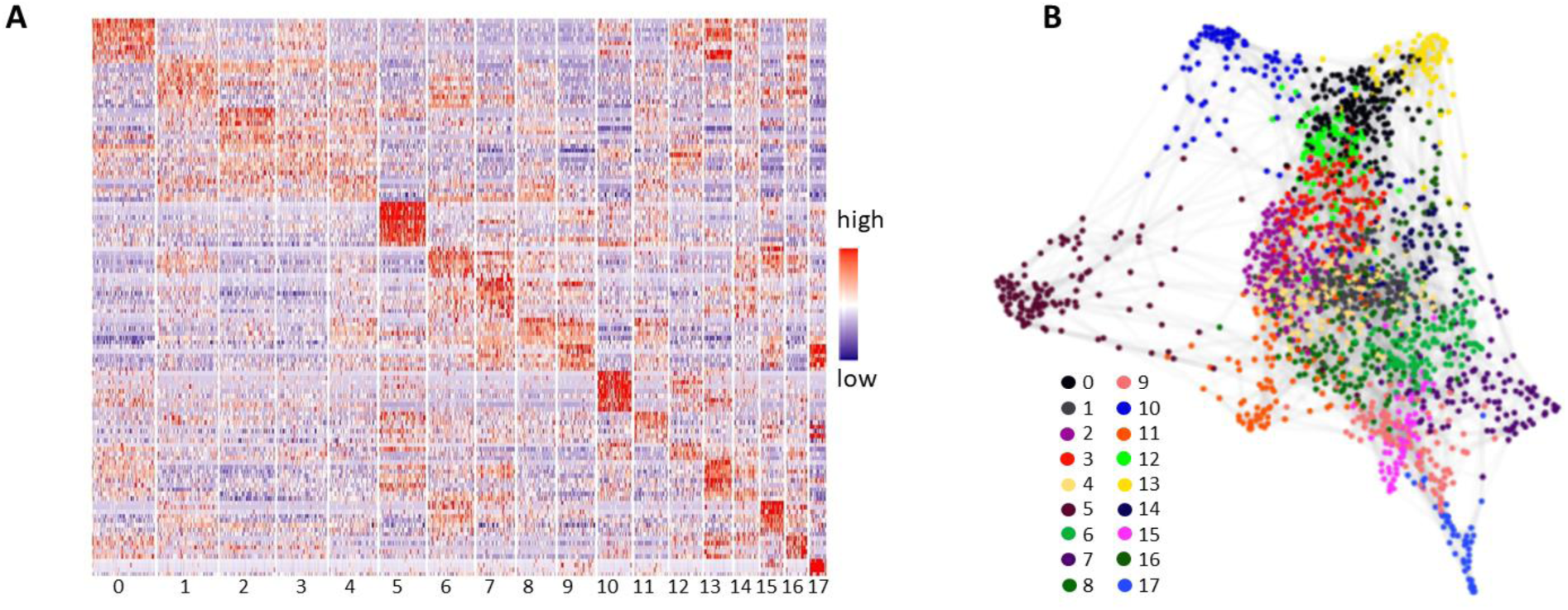
(A) Representative marker gene expression in each cluster. Only at most 10 representative marker genes are displayed for each cluster. (B) SPRING plot of the single cell transcriptome. Each cell is in the same color as they are in Figure 1B.

**Figure 2S.**
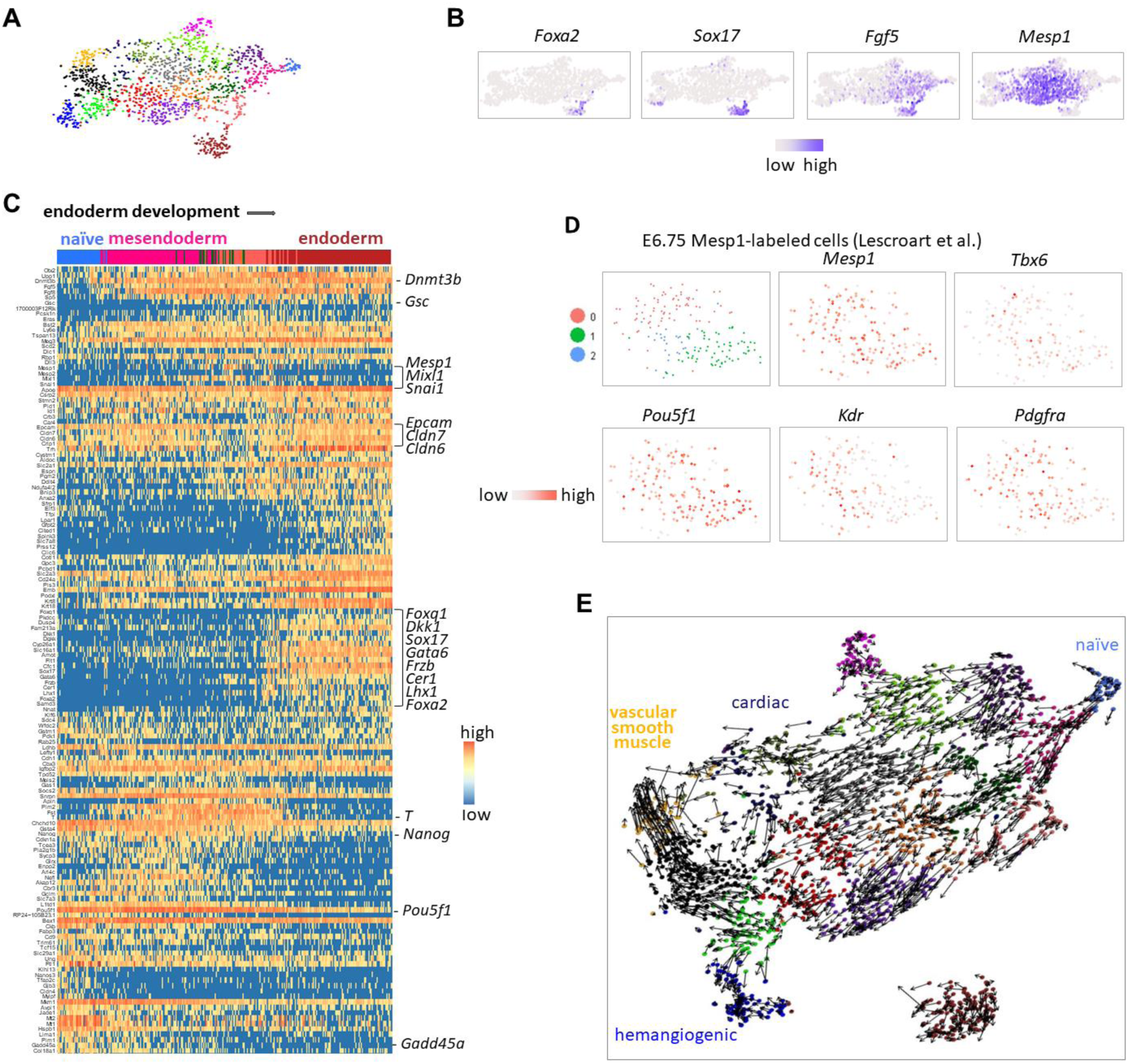
(A) Dynamics of the 163 most variable genes along the endoderm development route. The genes were clustered based on Pearson’s correlations. (B) t-SNE plot of the cells embedded with marker genes selected based on less strengency. The cells’ colors show their original identities in Figure 1B. (C) Gene expression in the new t-SNE plot. (D) Expression of indicated genes in E6.75 *Mesp1*-labeled cells (re-analysis of Lescroart et al.’s^8^ data). (E) RNA velocity estimation of the cells. The arrows indicate estimated differentiation trends of given cells. The colors of the cells is same to that in Figure 1B.

**Figure 3S.**
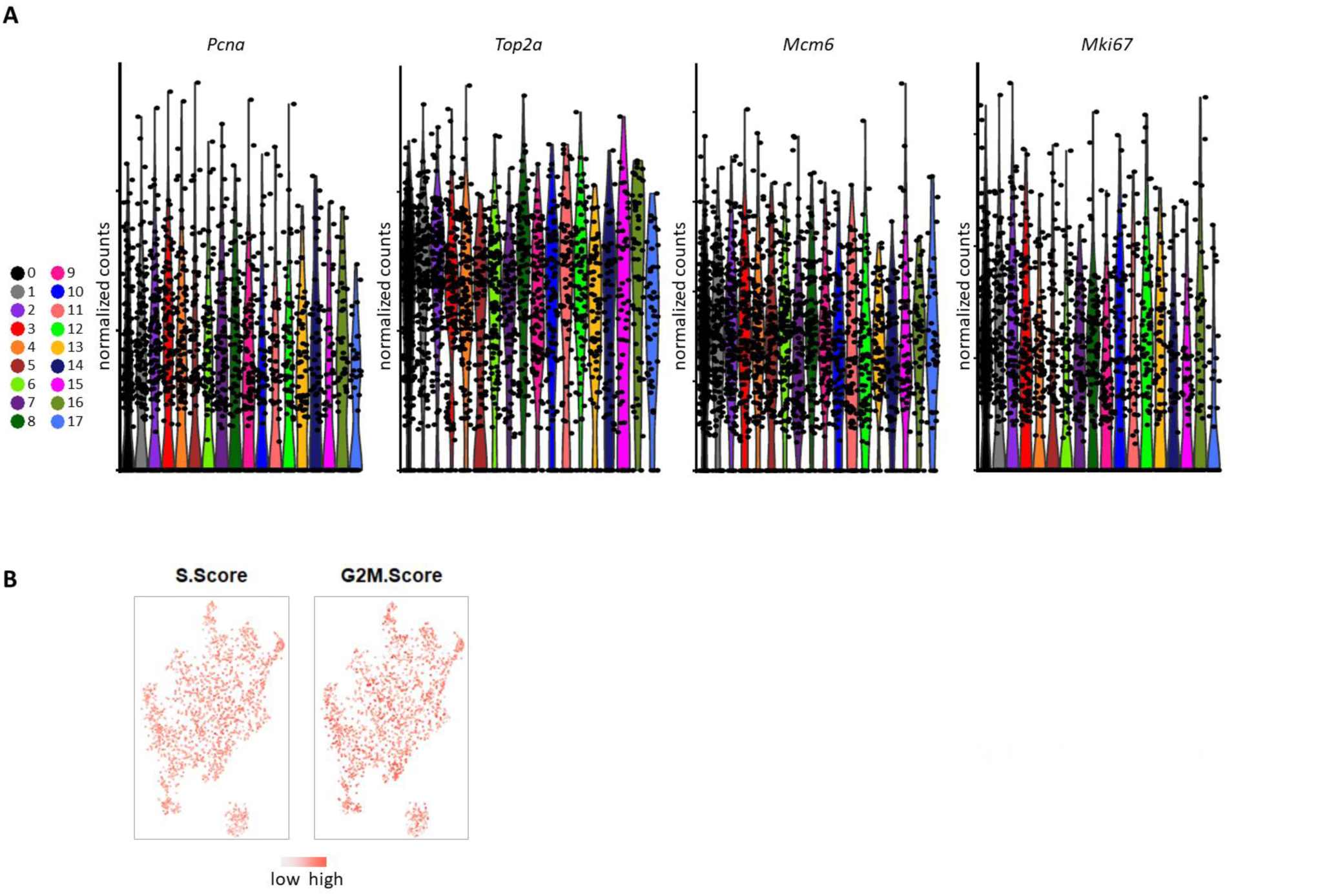
(A) Vln plot of cell cycle genes in different clusters. (B) Cell cycle scores of each cells.

**Figure 4S.**
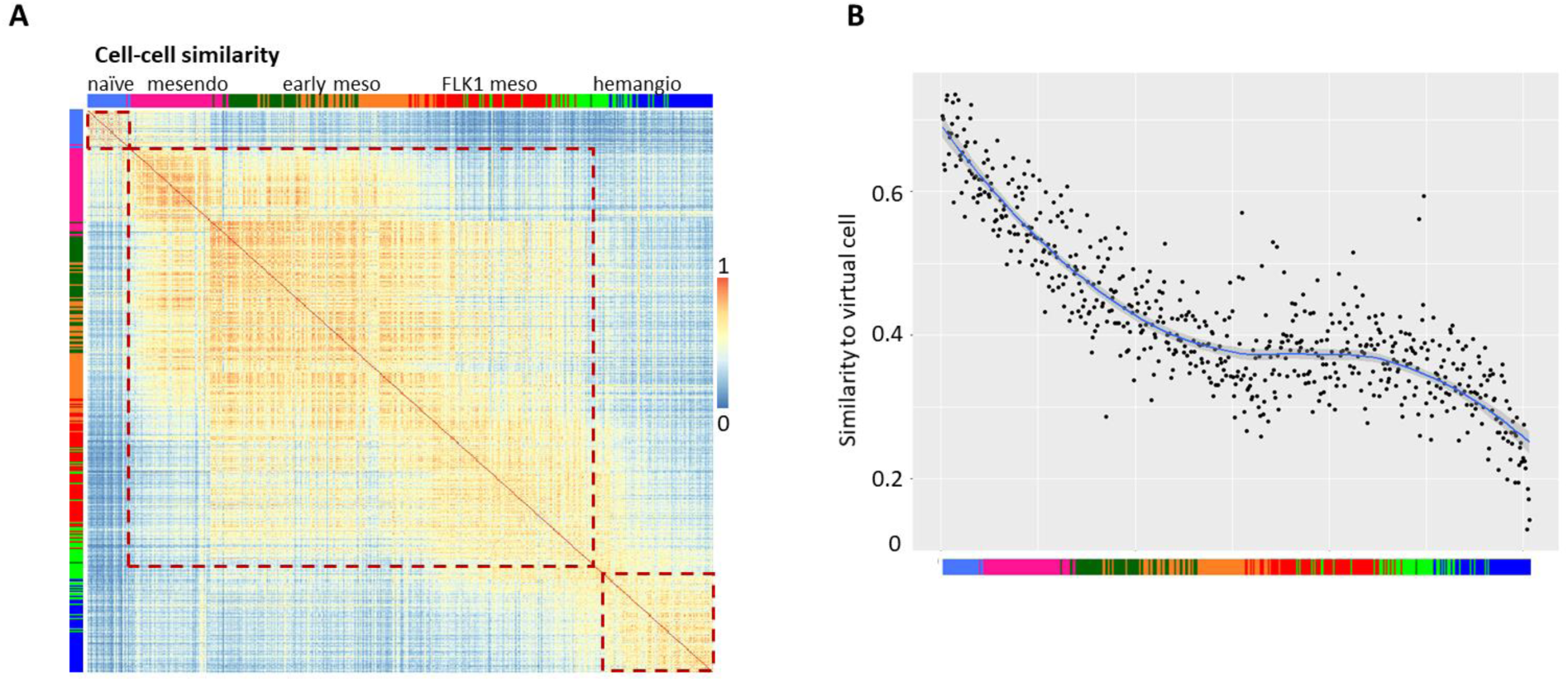
(A) Cell-cell Pearson’s correlations along the pseudo-time line based on expression of 427 most variable genes. The brown dash boxes mark the three relatively isolated naïve pluripotent population, ‘gastrulation’ stage, and hemangiogenic lineage. (B) The first 20 cells in the hemangiogenic lineage route were picked and their mean expression was treated as a virtual cell. Then each cell’s similarity in the development route to this virtual cell was indicated in the dot plot. Bars at the bottom show each cell’s original identity.

**Figure 5S.**
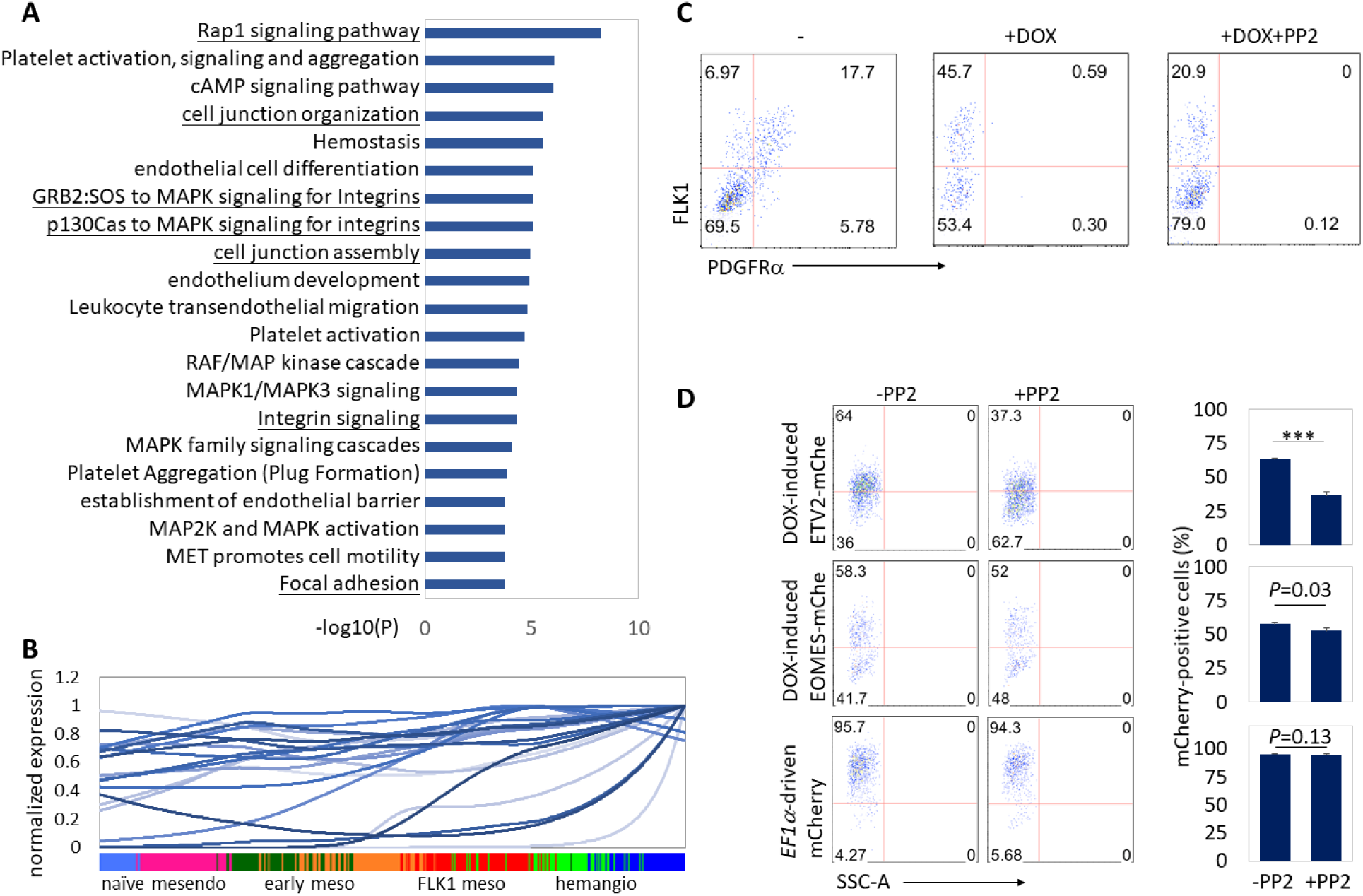
(A) Top 100 genes that are most required for *Tal1* expression and the top 100 most required for *Lmo2* expression have 66 overlap. These 66 genes were analyzed for functional enrichment. (B) Expression of 22 cell adhesion-related genes along the pseudo-time line of hemangiogenic lineage development. The expression values were smoothened for visualization. (C) A2lox ES cells with DOX-inducible exogenous *Etv2* expression were differentiated in serum medium for 3 days, EBs were then treated with or without the SRC kinase inhibitor PP2 (5μM) for another 24 hours in the presence of 2μg/mL of DOX. EBs without PP2 or DOX treatment were used as control. EBs were collected and analyzed using flow cytometry for surface marker examination. (D) A2lox ES cells with DOX-inducible ETV2-mCherry or EOMES-mCherry expression, or *EF1α*-driven constitutive mCherry expression were cultured in ES cell medium, treated with or without the SRC kinase inhibitor PP2 (5μM) in the presence of 0.5μg/mL of DOX for 1 days. Cells were collected and analyzed using flow cytometry for mCherry and FLK1 expression. The statistics summary is shown on the right. ***, P<0.001.

**Figure 6S.**
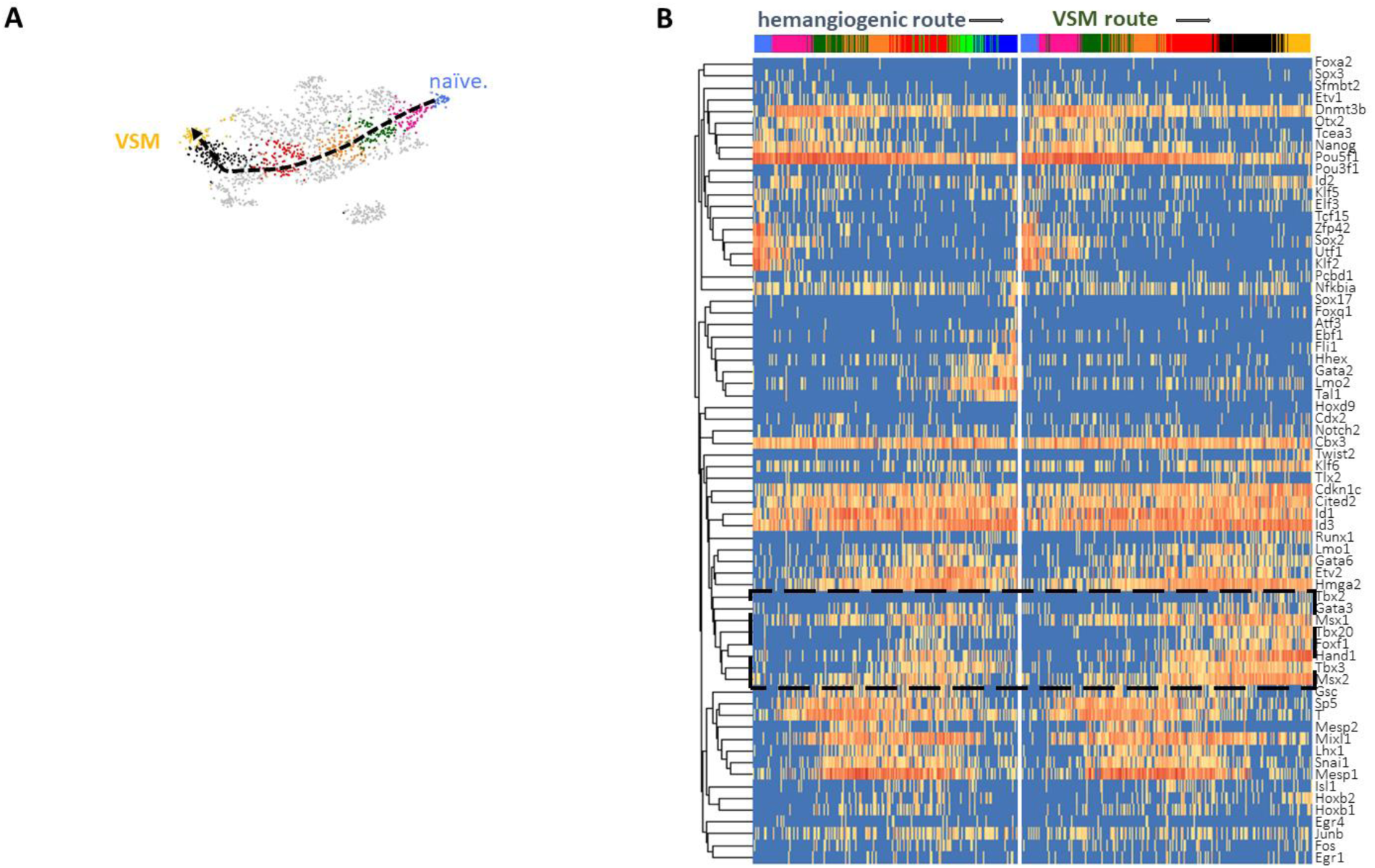
(A) Populations selected for VSM development route reconstitution. (B) Comparison of expression dynamics of transcription-related factors along VSM vs. hemangiogenic lineage development. The genes were clustered based on the Pearson’s correlations.

**Figure 7S.**
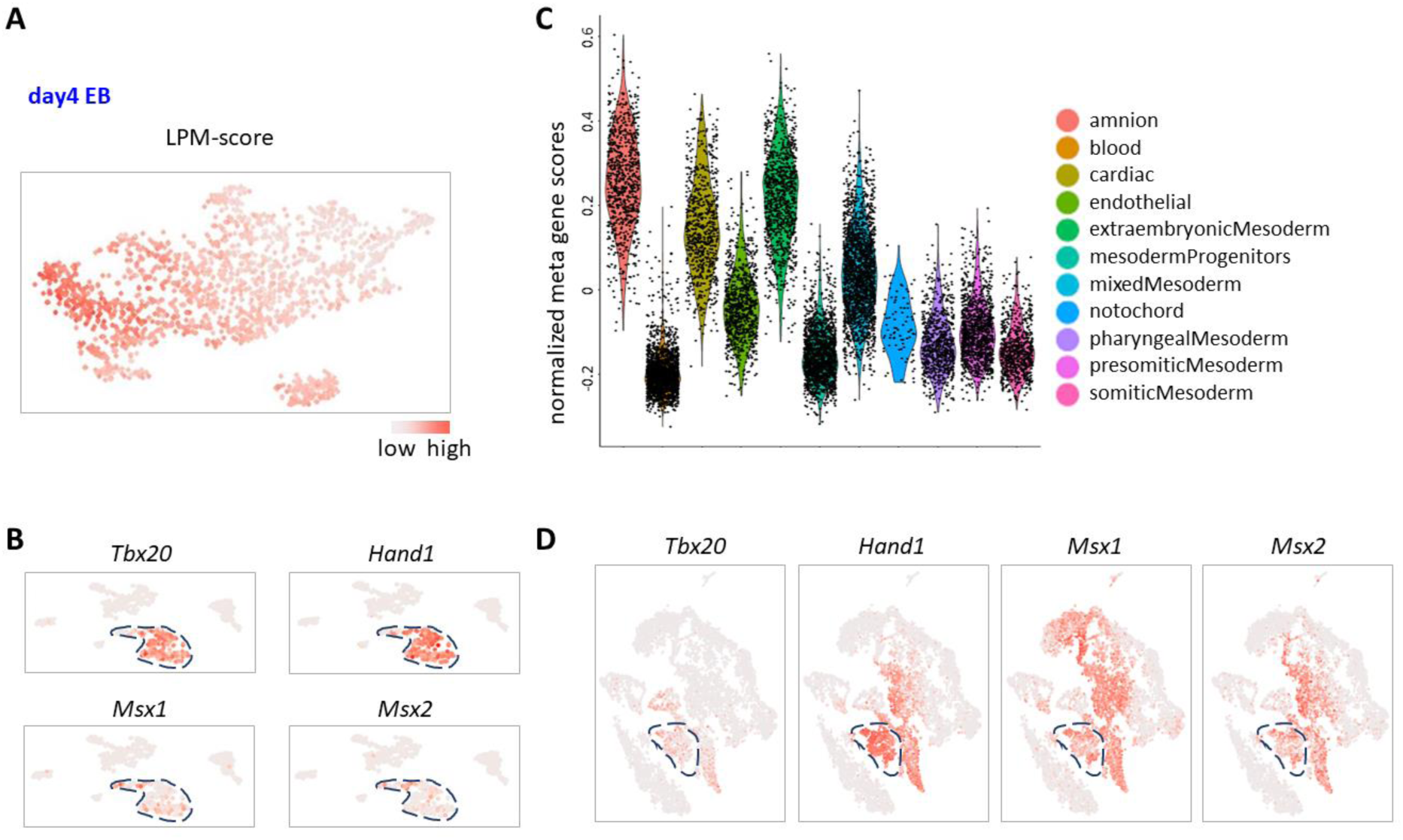
(A) LPM-score gene expression in day 4 EB cells. (B) Expression of indicated genes in E8.5 yolk sac cells. (C) Vln plot of FLK1-meata gene expression in different mesoderm tissues from E8.25 embryo. (D) Expression of indicated genes in E8.25 embryonic mesoderm cells.

